# Connecting genetic risk to disease endpoints through the human blood plasma proteome

**DOI:** 10.1101/086793

**Authors:** Karsten Suhre, Matthias Arnold, Aditya Bhagwat, Richard J. Cotton, Rudolf Engelke, Annika Laser, Johannes Raffler, Hina Sarwath, Gaurav Thareja, Robert Kirk DeLisle, Larry Gold, Marija Pezer, Gordan Lauc, Mohammed A. El-Din Selim, Dennis O. Mook-Kanamori, Eman K. Al-Dous, Yasmin A. Mohamoud, Joel Malek, Konstantin Strauch, Harald Grallert, Annette Peters, Gabi Kastenmuller, Christian Gieger, Johannes Graumann

## Abstract

Genome-wide association studies (GWAS) with intermediate phenotypes, like changes in metabolite and protein levels, provide functional evidence for mapping disease associations and translating them into clinical applications. However, although hundreds of genetic risk variants have been associated with complex disorders, the underlying molecular pathways often remain elusive. Associations with intermediate traits across multiple chromosome locations are key in establishing functional links between GWAS-identified risk-variants and disease endpoints. Here, we describe a GWAS performed with a highly multiplexed aptamer-based affinity proteomics platform. We quantified associations between protein level changes and gene variants in a German cohort and replicated this GWAS in an Arab/Asian cohort. We identified many independent, SNP-protein associations, which represent novel, inter-chromosomal links, related to autoimmune disorders, Alzheimer's disease, cardiovascular disease, cancer, and many other disease endpoints. We integrated this information into a genome-proteome network, and created an interactive web-tool for interrogations. Our results provide a basis for new approaches to pharmaceutical and diagnostic applications.

## INTRODUCTION

Genome-wide association studies (GWAS) with intermediate phenotypes, such as metabolite and protein levels, can reveal variations in protein abundances, enzymatic activities, interaction properties, or regulatory mechanisms, which can inform clinical applications^1–3^. Studies with aptamer-, immunoassay-, and mass-spectrometry-based proteomics (**Supplemental Table 1**) have shown that protein levels are strongly modulated by variations in the nearby *cis* regions of their encoding genes^4–9^. However, proteomics-based genetic association studies have been limited by small protein panels or cohort sizes, and thus, have not systematically addressed associations across multiple chromosome locations. These *trans*-associations are key in establishing functional links between different risk-variants, because they reveal the downstream effectors in the pathways between *cis-*encoded risk-variants and disease endpoints.

Here, we use a highly multiplexed, aptamer-based, affinity proteomics platform (SOMAscan^TM^)^10^ in a GWAS to quantify levels of 1124 proteins in blood plasma samples. The SOMAscan aptamers were generated to bind specifically to proteins implicated in numerous diseases and physiological processes and to target a broad range of secreted, intracellular, and extracellular proteins, which are detectable in blood plasma (**Supplemental Figure 1**, **Supplemental Data 1**). We investigate samples from 1000 individuals of the population based KORA (Cooperative Health Research in the Region of Augsburg) study ^1^,^11^ and replicate the results in 338 participants of the Qatar Metabolomics Study on Diabetes (QMDiab) with participants of Arab and Asian ethnicities ^12^.

We identify 539 independent, SNP-protein associations, which include novel inter-chromosomal (trans) links related to autoimmune disorders, Alzheimer’s disease, cardiovascular disease, cancer, and many other disease endpoints. Our study represents the outcome of over 1,100 new genome-wide association studies with blood circulating protein levels, many of which have never been reported before. We find that up to 60% of the naturally occurring variance in the blood plasma levels of essential proteins can be explained by two or more independent variants of a single gene located on another chromosome, and identify strong genetic associations with intermediate traits related to proteins involved in pathways to complex disorders.

## RESULTS

### A genome-wide association study with 1124 proteins

We used linear additive genetic regression models, adjusted for relevant covariates, to analyse 509,946 common autosomal single nucleotide polymorphisms (SNP_s_) for genome-wide associations with 1124 protein levels measured in 1000 blood samples from the KORA study (**see Methods**). We grouped association signals with correlated SNP_s_ (r^2^>0.1) into single protein quantitative trait loci (pQTLs) and identified 539 pQTLs (284 unique proteins, 451 independent SNPs) with a conservative Bonferroni level of significance (p<8.72×10^-11^ = 0.05/509,946/1124). For 21% of the assayed proteins we detected a cis-pQTL at a more liberal significance cut-off of p<5×10^-8^. The full list is available as **Supplemental Data 2**. We then fine-mapped our associations with variants imputed from the 1000-Genomes project database. Next, we attempted replication of 462 associations that had a suitable genotyped tag SNPs (r^2^>0.8) in samples from 338 participants of the QMDiab study^12^. Of those, 234 associations were replicated at a Bonferroni level of significance (p<1.08×10^-4^ = 0.05/462), and an additional 150 showed nominal significance (p<0.05) in the replication sample. We observed directional consistency between primary and replication sample for 215 of the 234 replicated SNPs. Discrepancy for the remaining 19 SNPs may be explained by changes in major and minor allele coding between the two cohorts and ethnic differences. What is more, out of 208 associations that had 95% replication power (determined by sampling), 171 (82.2%) were replicated and 198 (95.2%) were nominally significant. Moreover, we replicated several associations previously reported in aptamer-, immunoassay-, and mass-spectrometry-based studies, which demonstrated concordance among these technologies (**Supplemental Table 1, Supplemental Table 2**, **Supplemental Table 3**, **Supplemental Table 4**, **Supplemental Table 5**).

Next, we identified and annotated putative causative and disease-relevant variants. We used the SNiPA web-tool^13^ to retrieve variant-specific annotations for all SNPs from the 1000-Genomes project in linkage disequilibrium (LD) with an identified pQTL (LD r^2^>0.8). The annotations included primary effect predictions, SNPs in experimentally identified regulatory elements (ENCODE), expression QTLs (eQTLs), and disease associated variants. Of 384 annotated *cis-*pQTLs, 228 had a variant in the gene coding region, whereof 74 were protein-changing. Eighty-eight had a variant in a regulatory element, and 179 had an eQTL that matched the associated protein. We complemented the pQTL annotation with 122 overlapping methylation QTLs (meQTLs) (**Supplemental Data 2**) and 14 overlapping metabolic QTLs (mQTLs) (**Supplemental Table 6**). With the GWAS catalogue as a reference, and by including publicly available summary statistics data from 16 large disease GWAS consortia, we identified 83 GWAS associations for 42 unique disease endpoints that overlapped with 57 pQTL loci (**Supplemental Table 7**). With the Ingenuity Pathway Analysis database (IPA, Qiagen Inc.), we annotated 50 proteins as clinical or pharmaceutical biomarkers (**Supplemental Table 8**) and 43 as drug targets (**Supplemental Table 9**), and all of these had at least one replicated pQTL.

We used Gaussian graphical modelling (GGM) to connect 1092 proteins through 3943 Bonferroni-significant partial protein-protein correlation edges (**Supplemental Data 3**). Then, we added all 451 genetic pQTL variants as nodes, and connected them to the protein network through the 539 SNP-protein associations (Figure 1). This network is freely available online, and it can be navigated via an interactive web interface. Overall, we found that given our study power the blood plasma levels of over 20% of all assayed proteins were under substantial genetic control, and in some cases, this control resulted in nearly total protein ablation (Figure 2). Additionally, a wide-spread feature was the convergence of multiple association signals to impact the levels of key proteins of biomedical and pharmaceutical interest (**Supplemental Figure 2**).

**Figure 1:**
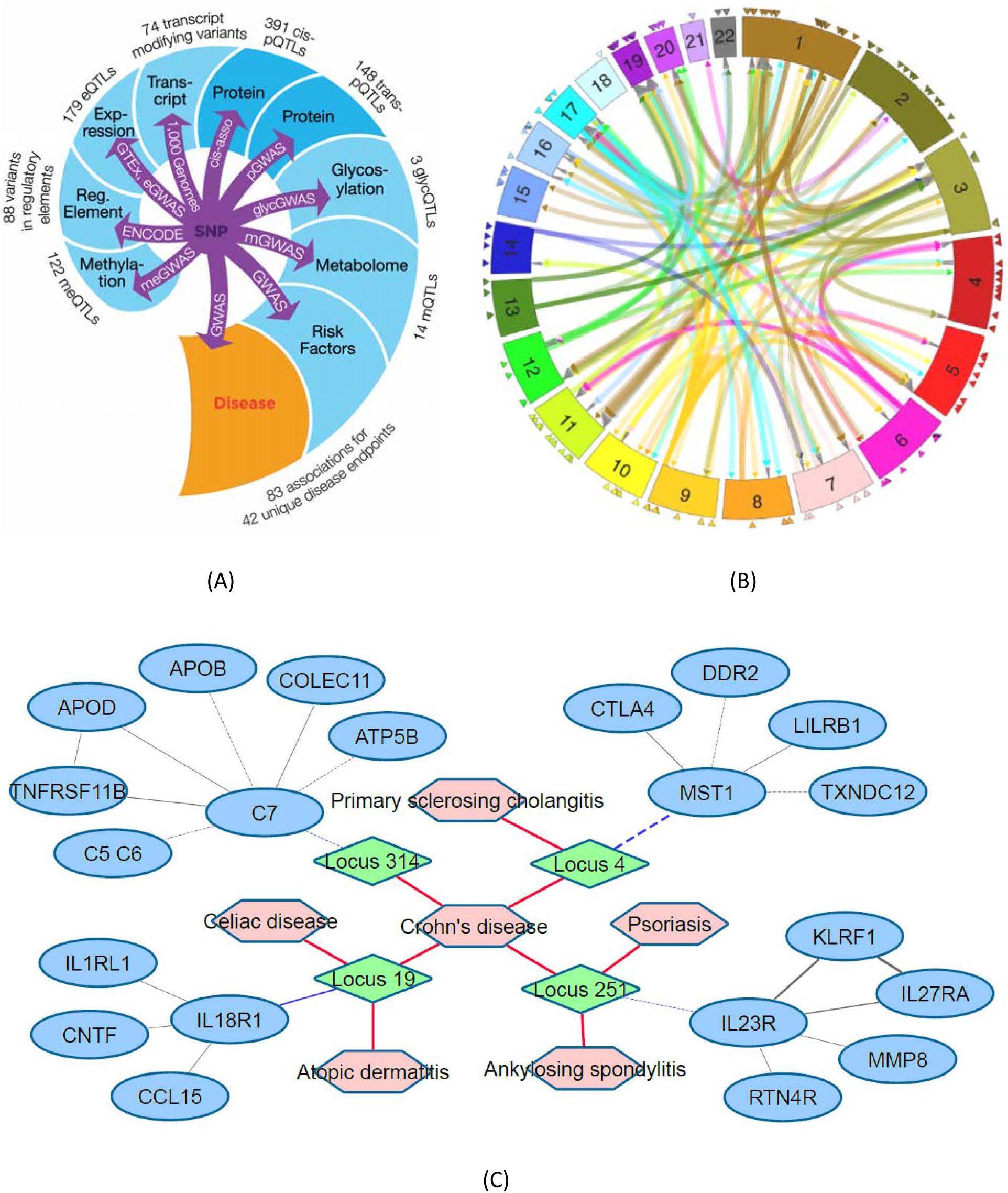
The genome-proteome-disease network. (A) Data sources integrated into the network, indicating the number and type of the overlapping associations, from the SNP to the disease endpoint; all associations are freely accessible at http://proteomics.gwas.eu; (B) Circular plot of all *cis-* and *trans*-associations, *cis*-pQTLs are indicated by triangles, *trans*-pQTLs connect associated variant locations and *trans*-encoded protein locations, an interactive version of this circular plot constitutes an entry point to query the integrated web-server; (C) Example of a genome-proteome-disease sub-network obtained from the server for a query using the search word *“Crohn’s Disease”*; network elements are disease traits (salmon hexagons), pQTL loci (green diamonds), protein levels (blue ovals); nodes are connected by genetic associations, partial correlations, and disease GWAS associations; This example (edited here for clarity) revealed four risk loci that associated with plasma levels of C7, MST, IL23R, and IL18R, respectively; These four proteins all play a major role in auto-immune disorders; Partial correlations between neighbouring proteins reveal pathways that may be involved in the aetiology of Crohn’s disease; Similar networks can be retrieved starting with a query using any of the 539 pQTLs, 1124 proteins and 42 unique co-associated disease endpoints; All items are interactively linked to association data from the discovery and the replication study, regional association plots based on imputed variants, locus annotations including co-associated eQTL-, meQTL-, mQTL-, regulatory-, coding-, and disease risk-variants, and link-outs to relevant protein databases, original data sources, and primary publications. The links in this network reflect the outcome of many natural experiments, represented by genetic variations observed in the genomes of hundreds of individuals from the general population and probed by deep proteomics phenotyping using over 1000 aptamers. The example of the network shown in (C) refers to Chrohn’s Disease and illustrates that four genetic loci identified by GWAS link proteins from the complement system and cytokines implicated in inflammatory processes to the disease.

**Figure 2:**
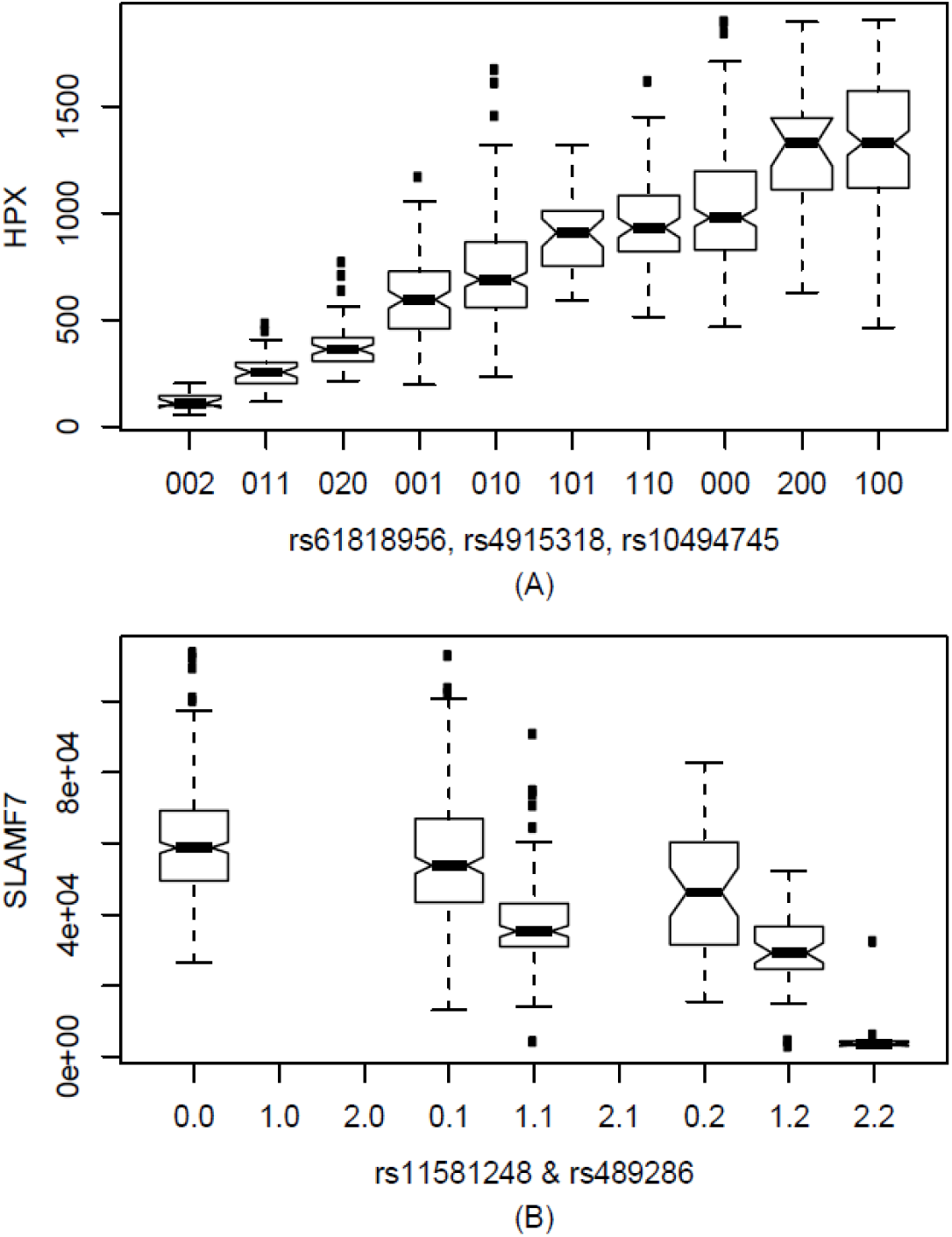
Examples of protein levels that are determined by multiple independent genetic variants. Box plots of protein levels of Hemopexin HPX (A) and SLAMF7 (B) as a function of genotype. The number of minor alleles of the respective genetic variant is given; for instance, in (A), “002” refers to individuals that are homozygous for the major alleles of rs61818956 and rs4915318 and for the minor allele of rs10494745, and in (B), “0.2” refers to homozygotes of the major allele of rs11581248 and the minor allele of rs489286. Only variant combinations that were observed in the study population are shown in the case of HPX. SNPs rs61818956, rs4915318, and rs10494745 are located in *trans* in the complement factor H-related 2/4 (CFHR2/CFHR4) gene locus. Further examples are shown in **Supplemental Figure 2**.

### Trans-associations

*Trans-*associations are exceptionally valuable for identifying new pathways. These associations establish causal links between proteins encoded at the GWAS loci and the blood levels of one or several *trans-*encoded proteins. We identified 148 *trans-*pQTLs and replicated 55. Forty-nine *trans-*pQTLs had 95% replication power, we replicated 38 of these. Six replicated *trans-*pQTLs had an additional replicated *cis-*association, two had two replicated *trans-*associations and one had three replicated *trans-*associations. Three independent SNPs at the haptoglobin (HP) locus had together four pQTLs, and two proteins had replicated *trans-*associations at two distinct chromosome locations (Table 1).

**Table 1:**
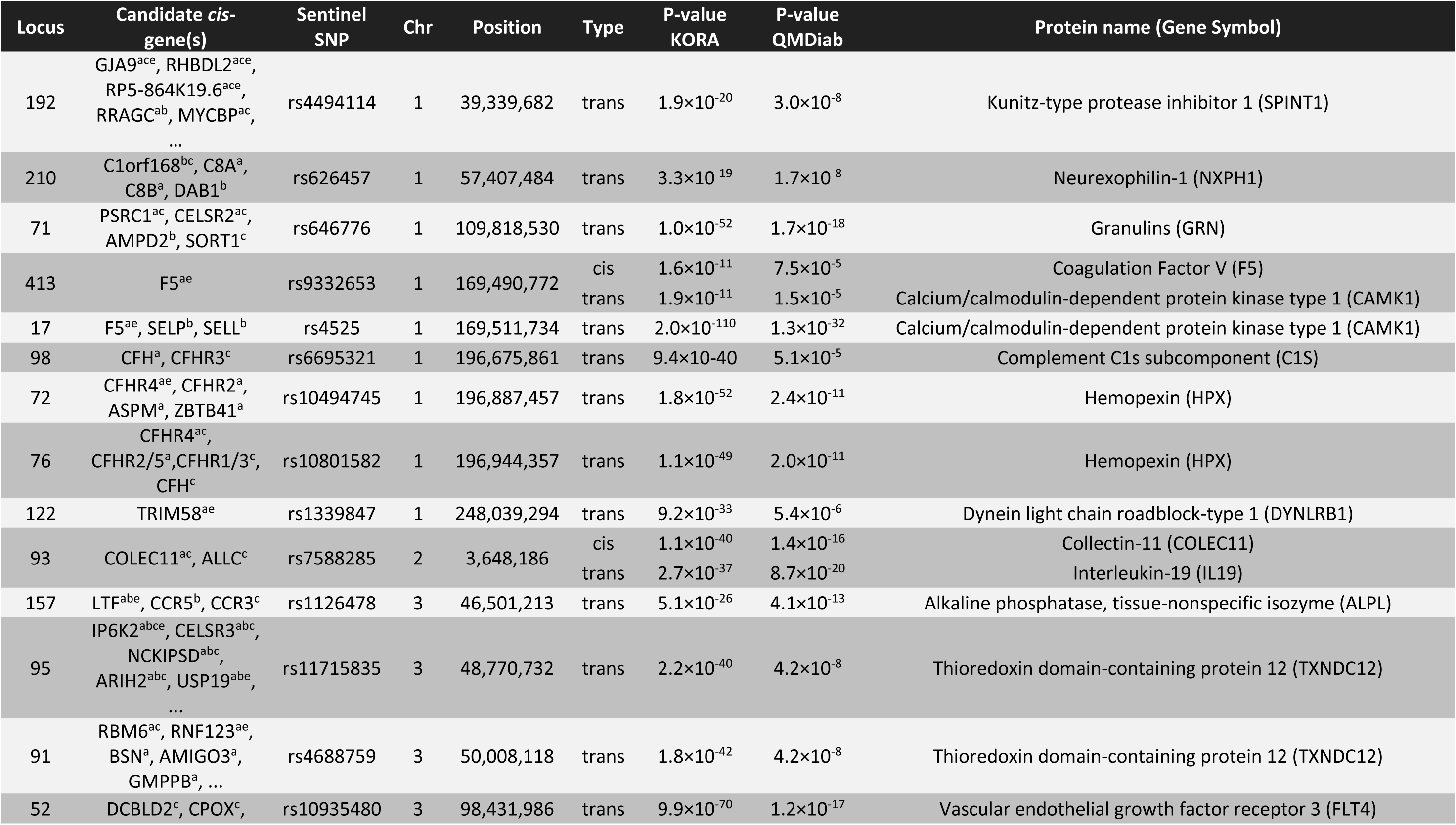
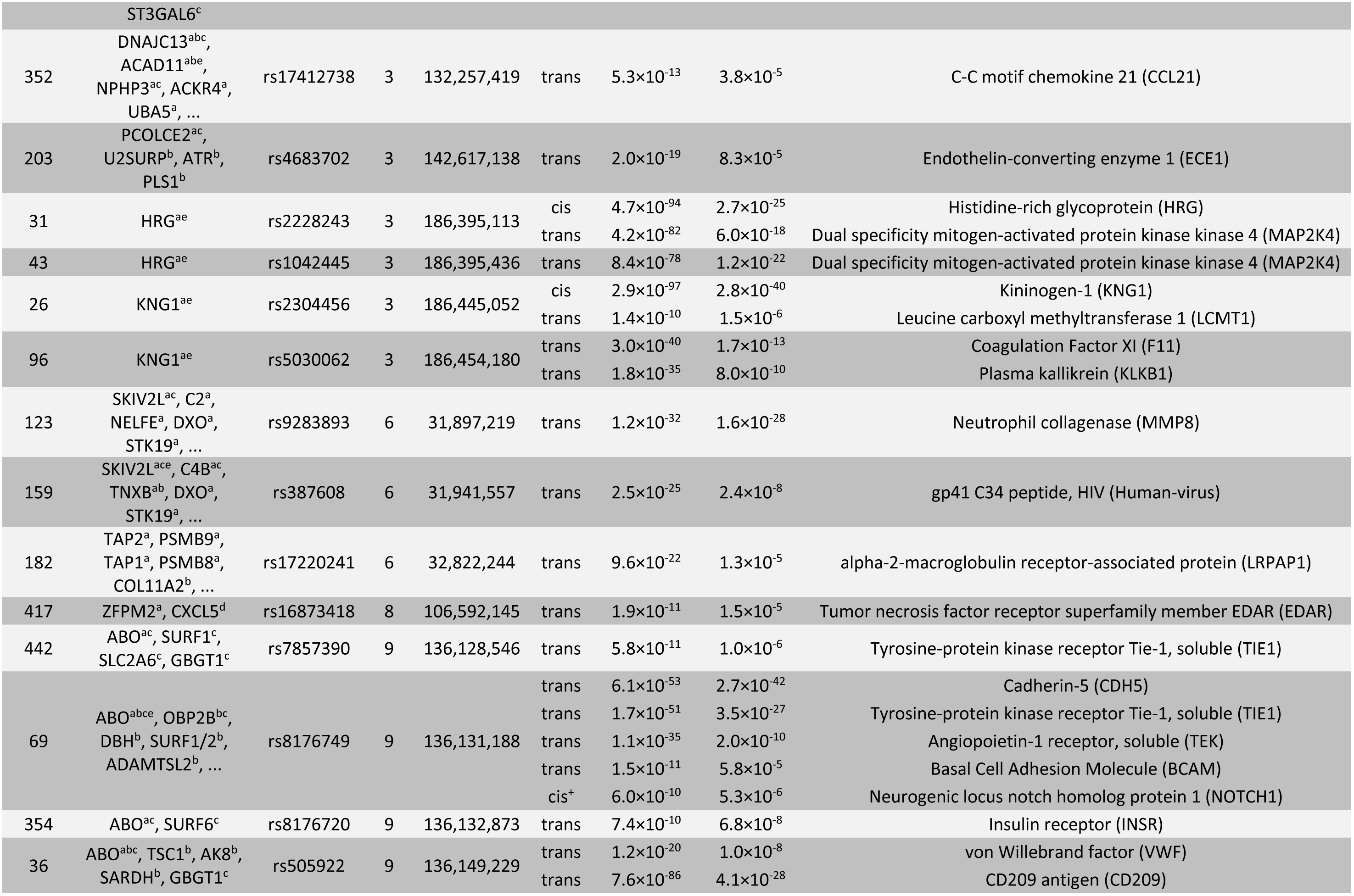
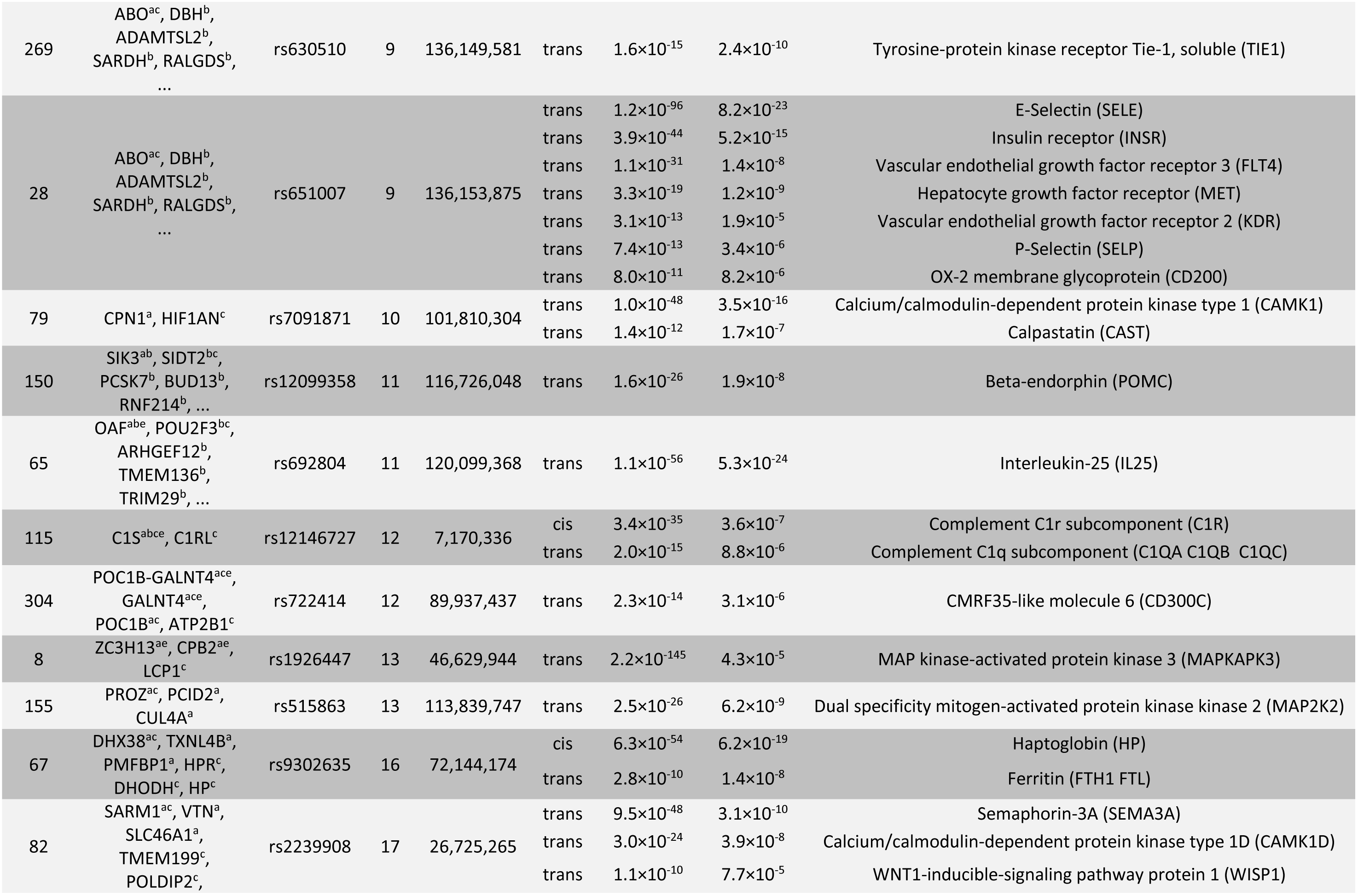
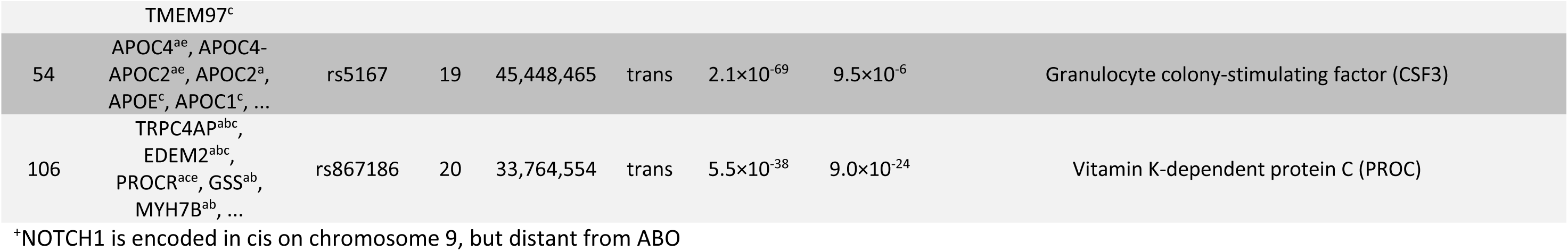
List of replicated *trans-* pQTLs. Loci that comprise at least one replicated *trans-* association. Loci are referenced in this study by numbers ranging from 1 to 451 (strongest to weakest) and sorted here by chromosome position. P-values are for the association with inverse-normal scaled protein levels; see **Supplemental Data 2** for full data of all 539 SNP-protein associations at 451 loci, including statistics for association with alternatively raw- and log-normal-scaled protein levels and estimated replication power. Candidate genes for the protein associations were annotated by considering the following criteria: a variant in LD with the sentinel SNP (r2>0.8) is located in the gene transcript (a), a variant hits a regulatory element of that gene (b), a variant is a *cis-* eQTL (c), a variant is a *trans-* eQTL (d), a variant is protein changing (e). The list of candidate genes in this table is limited to the five most plausible candidate genes for each locus; the full list is available online and in **Supplemental Data 2**. Every *trans-* pQTL implies the existence of a functional and causal link between a *cis-* encoded candidate gene and the target protein(s).

In this study, the pleiotropic ABO blood group gene exhibited the most promiscuous protein association signal. This locus displayed six independent genetic variants that were associated with 14 different proteins through replicated *trans-* pQTLs; three of these associations had been published previously (VWF, SELE, SELP, **Supplemental Table 2**). The other eleven associations were new, to our knowledge (BCAM, CD200, CD209, CDH5, FLT4, INSR, KDR, MET, NOTCH1, TEK, TIE1).

Genetic variance in ABO has been associated with coronary artery disease and stroke ^14^ and diabetes ^15^ (**Supplemental Figure 3**). The non-O blood group is one of the most important genetic risk factors for venous thromboembolism^16^, pancreatic cancer^17^, and susceptibility to infectious diseases ^18^. However, we lack a full understanding of the proteins and pathways involved in the pathogenic effects of these genetic variants. Based on the pQTLs reported here, we found support for the following new hypotheses: (1) The association between ABO and the Insulin receptor (INSR) reflects the well-established associations between the ABO locus and diabetes and the insulin receptor and diabetes ^15^. The association between ABO and the Insulin receptor (INSR) suggest that INSR-mediated insulin signaling may be involved in the ABO-diabetes association. (2) rs651007 associated here with *P-selectin* (SELP). SELP-positive platelets were previously reported to be associated with blood pressure ^19^, and rs651007 was associated in a GWAS with angiotensin converting enzyme (ACE) activity. This variant was further identified as an mQTL for a number of dipeptides in blood ^20^, which may be produced by the dipeptidase ACE ^21^. These observations suggest a potential role for the *SELP* pQTL in the GWAS association between ABO and cardiovascular disease. (3) Several of the 14 proteins associated here with ABO have been shown to interact or form complexes in relation with angiogenesis and vascular maturation processes: *Angiopoietin-1 receptor, soluble* (TEK) plays a role in embryonic vascular development and phosphorylates *Tyrosine kinase with immunoglobulin-like and EGF-like domains 1* (TIE1). TIE1 overexpression in endothelial cells upregulates selectin E (SELE) ^22^. Strain induced angiogenesis is mediated in part through a Notch-dependent, Ang1/Tie2 signalling pathway that implicates NOTCH1 and TIE1 ^23^. *Vascular endothelial (VE)-cadherin* (CDH5) is required for normal development of the vasculature in the embryo and for angiogenesis in the adult, and it is associated with *VE growth factor (VEGF) receptor-2* (KDR) on the exposure of endothelial cells to VEGF ^24^. Heterodimers of KDR and *Vascular endothelial growth factor receptor 3* (FLT4) positively regulate angiogenic sprouting ^25^. VEGF directly and negatively regulates tumor cell invasion through enhanced recruitment of the *Protein tyrosine phosphatase 1B* (PTP1B) to a *hepatocyte growth factor receptor* (MET) - KDR hetero-complex ^26^. VEGF also synergistically increased tumor necrosis factor-alpha-induced E-selectin mRNA and shedding of soluble E-selectin. Synergistic upregulation of E-selectin expression by VEGF is mediated via KDR and calcineurin signaling ^27^. These observations suggest that genetic variance in ABO has major effects on a network comprising a number of proteins involved in cell adhesion, angiogenesis and neo-vascularization processes. Consequently, the here reported *trans-*pQTLs indicate novel pathways that may be involved in ABO-mediated cancer susceptibility in addition to regulating vascular metabolism ^28^.

### Post-translational modifications

To follow up on the many hypotheses generated by the pQTLs reported here, expert knowledge and experimentation will be required. Given the fact that several of the proteins identified in our pQTLs were glycoproteins, we performed plasma protein glycoprofiling (see **Online Methods**) to investigate whether our pQTLs were involved in post-translational protein modifications.

**Figure 3:**
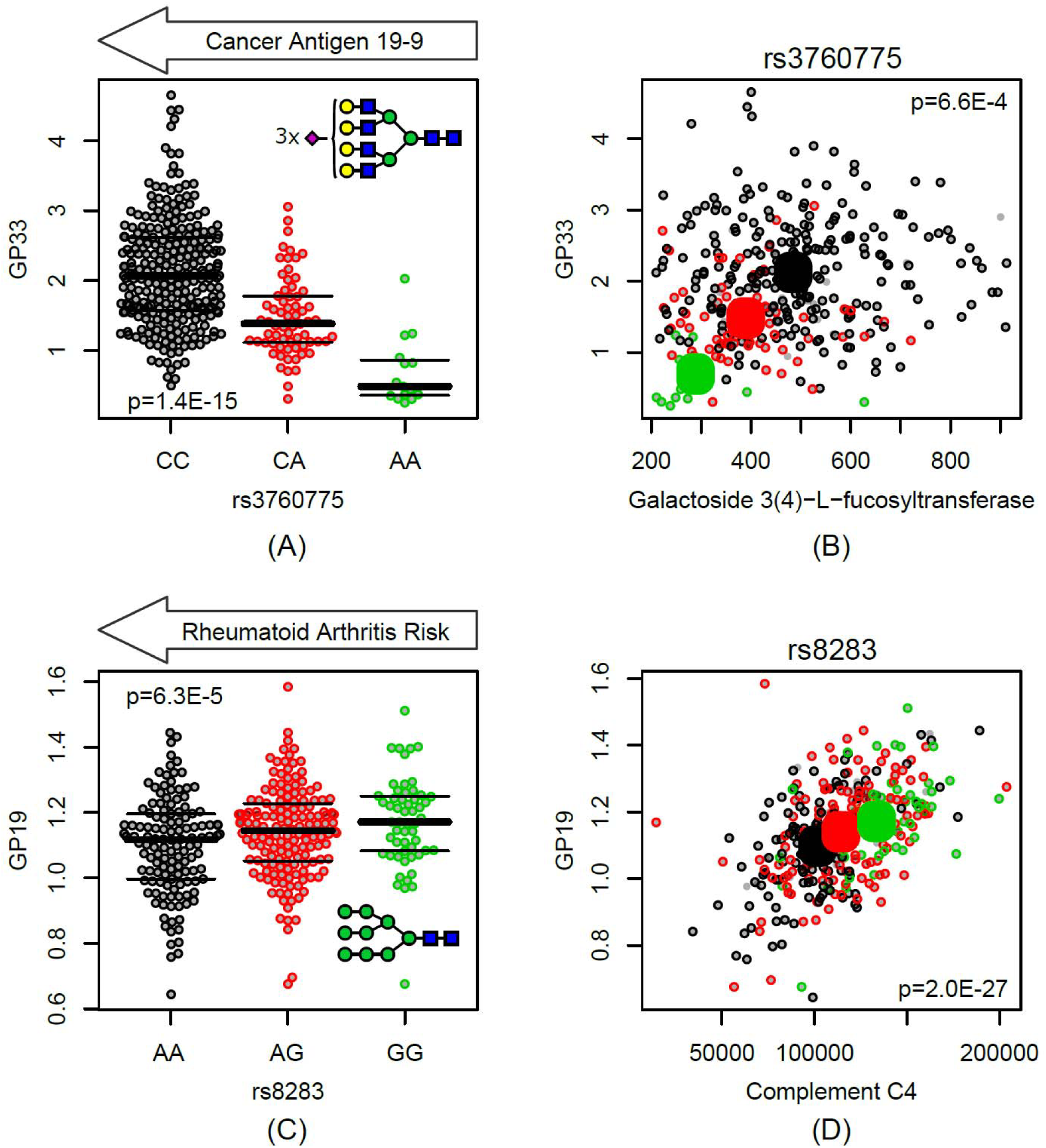
Genotype-dependent co-associations of the plasma proteome and the plasma N-glycome. Bee swarm plots of total plasma N-glycans GP19 and GP33 (% of total N-glycan content) as a function of rs3760775 and rs8283 genotype, respectively (A&C), see inset for glycan structure, Blue squares: N-acetylglucosamine, green circles: mannose, yellow circles: galactose, purple diamonds: N-acetylneuraminic acid; Scatter plots of total plasma N-glycans GP19 and GP33 as a function of *Complement factor 4* (C4) and *Galactoside 3(4)-L-fucosyltransferase* (FUT3) (raw data), respectively (B&D), coloured by genotype, black: major allele homozygotes, red: heterozygotes, green: minor allele homozygotes, large circles indicate means by genotype; p-values are for the association of glycans with genotype (A,C) and of glycans with protein levels (B,D); the major allele variant of SNP rs3760775 was reported to be associated with the cancer antigen 19-9 and that of SNP rs8283 with increased risk of rheumatoid arthritis.

For instance, SNP rs3760775, located near the *FUT3* gene, was associated with plasma levels of the corresponding galactoside3(4)-L-fucosyltransferase protein. We found that this same variant was strongly associated with the N-glycan GP33 (p=1.4×10^-15^) (see Figure 3A for GP33 structure). In a previous GWAS, SNP rs3760775 was reported to be associated with the glycan antigen, CA19-9 ^29^, a widely used cancer biomarker, which is present on multiple proteins ^30^. Nearly 20 years ago, studies demonstrated a similar association between CA19-9 and the *FUT3* gene dosage ^31,32^. This previously reported association between variance in the FUT3 locus and the expression of cancer antigen, CA19-9, and our observation of a similar association with GP33, suggest that both glycans may be involved in a same pathway.

Another glycoprofile investigation began with the strong, genotype-dependent correlation between complement factor C4 (C4) and the N-glycan, GP19 (p=2.0×10^-27^) (Figure 3D). We also found that SNP rs8283 was associated with both C4 and GP19. GP19 is composed of 9 mannose moieties (M9 glycan structure), and it was previously reported to be attached to C4 ^33^. Moreover, M9 glycans are the principal target ligand for mannose binding protein (MBL), which binds to C4 ^34^, and subsequently, activates the complement system through the lectin pathway. MBL has been implicated in the pathology of rheumatoid arthritis (RA) in many studies, including a recent meta-analysis, which confirmed the association between functional MBL variants and RA risk ^35^. The rs8283 variant was also associated with RA (p=3.8×10^-51^)^36^, but is not in linkage disequilibrium with the top reported RA-risk variants in the HLA region. Hence, SNP rs8283 appears to be an independent signal, possibly mediated through MBL binding to C4 glycans.

### Biomedical relevance

A major challenge in conducting a disease GWAS is the difficulty in identifying causative variants in the pathophysiological pathways that lead to the observed clinical manifestations. Generally, hypotheses about the identity of the disease-causing genes are based on biological arguments, such as the presence of SNPs in the regulatory or coding regions of functionally plausible genes, which may be supported by co-associated eQTLs. However, a much stronger argument is provided by the co-association with a pQTL, which constitutes firm experimental evidence that the blood levels of disease-associated proteins vary in response to changes in the genome. Moreover, partial correlations to functionally related proteins, as reported here, may further substantiate hypotheses generated from pQTLs (see example in Figure 1C). In the following sections, we show how pQTLs identified in this study reveal new insights into multiple disease-associated pathways identified in previous GWAS.

### Auto-immune disorders

Ankylosing spondylitis (AS) is a common cause of inflammatory arthritis, and it affects one in 200 Europeans. Evans et al.^37^ identified two AS-risk variants in the endoplasmic reticulum aminopeptidase 1 (*ERAP1*) gene; they reported that the major allele of rs30187 and the minor allele of rs10050860 were protective. ERAP1 is involved in trimming peptides prior to HLA class I presentation; it has recently attracted attention as a drug target for auto-immune disorders^38^. Several studies showed that ERAP1 was present in blood; it was localized to exosome-like vesicles and present in the extracellular space^39^. Here, we found that the two identified AS-risk variants were associated, in an additive manner, with increasing levels of circulating ERAP1 protein (Figure 4A). Similarly, mRNA sequences isolated from lymphoblastoid cells showed that ERAP1 mRNA expression also increased with increasing numbers of AS-risk alleles (Figure 4B). Previous work concluded that the association between ERAP1 and AS was mainly driven by genetic differences in how ERAP1 enzymatic activity shaped the HLA-B27 peptidome ^40,41^. Our findings suggest that the auto-immunogenic effects of different ERAP1 protein variants may be modulated by genotype-dependent regulation of protein expression. This observation may have broader implications for treatment approaches to autoimmune disorders that depend on ERAP1 antigen processing.

**Figure 4:**
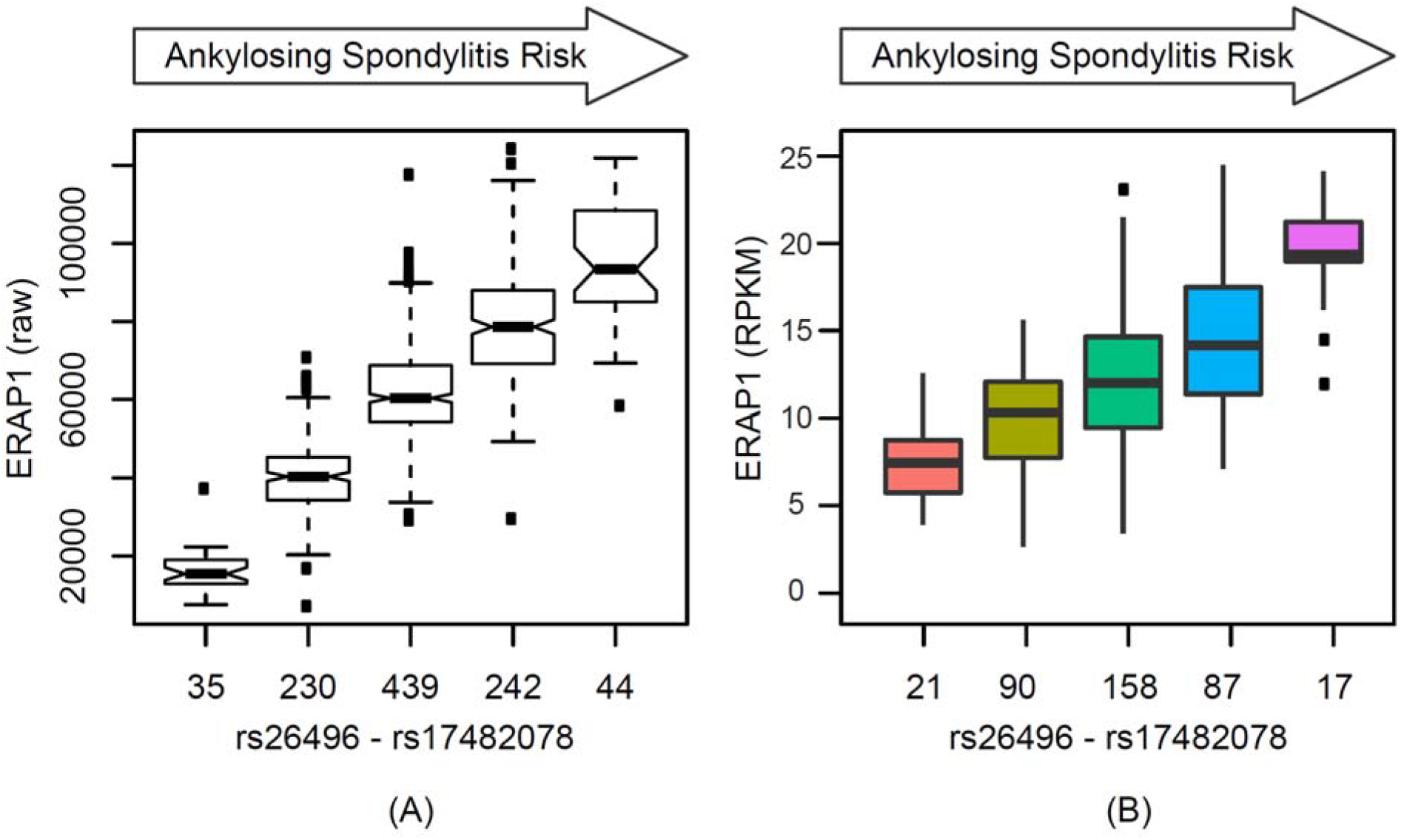
Protein and mRNA expression levels of endoplasmic reticulum aminopeptidase 1 (ERAP1) as a function of two Ankylosing spondylitis (AS)-risk alleles. Boxplots of ERAP1 blood circulating protein levels (A) and ERAP1 mRNA expression levels observed in lymphoblastoid cells (B) as a function of the sum of AS-risk alleles (minor allele of rs26496, r^2^=0.46 with rs30187; major allele of rs17482078, r^2^=0.96 with rs10050860); the number of individuals/cell lines with the respective genotype is indicated below the x-axis.

### Complement system and haem clearance

We identified two independent variants (rs10494745 and rs10801582), located in the complement factor H-related 2/4 (CFHR2/CFHR4) gene locus. These variants were associated in *trans* with haemopexin (HPX) protein levels (Figure 2A). Lower CFHR4 expression levels were associated with lower HPX protein levels. HPX binds haem with high affinity and transports it from the plasma to the liver, which prevents the accumulation of oxidative species. The rs10494745 variant is a G→E amino acid substitution in CFHR4. It is an eQTL for CFHR4 expression in liver, as it tags the GTEx rs4915318 variant (GTEx, p=1.6×10^-7^). Imputed data revealed a third, strong, and independent signal on SNP rs61818956 (p=1.13×10^-74^), which is located in an intron in the CFHR2/CFHR4 locus. Conditional analysis showed that all three variants were statistically independent, and together, they explained a surprising 61% of the observed variance in a key protein responsible for oxidative stress reduction. Previous studies reported that CFHR4 interacts with complement component 3 (C3) ^42^, and in turn, C3 interacts with haem ^43^. Those findings suggest a plausible *trans-*acting pathway that links CFHR4 and HPX. That observation may have important consequences on our understanding of the pathologies involved with the classical and alternative complement activation pathways.

### Alzheimer’s disease (AD) and mRNA splicing

Our findings on the major AD-risk variant, rs4420638, may generate particular medical interest. This variant displayed a *cis-*association with increased levels of apolipoprotein E (isoform E2) (APOE) and a concordant *trans-*association with decreased levels of small nuclear ribonucleoprotein F (SNRPF). This dual association was further supported by the finding that the ratio between APOE and SNRPF strengthened the association with rs4420638 by 16 orders of magnitude (p-gain statistic ^44^). Although this association lacked sufficient replication power, both APOE and SNRPF associations remained nominally significant in the QMDiab (p<0.012), with corresponding trends. SNRPF is a core component of U1, U2, U4, and U5 small nuclear ribonucleoproteins (snRNPs), which are the building blocks of the spliceosome. Recently, a knock-down of U1-70K or inhibition of U1 snRNP components was shown to increase the levels of amyloid precursor protein ^45^. An association between this major AD-risk variant and a protein of the spliceosomal machinery has not been reported previously. Taken together, these observations support the implication that protein splicing may be an important factor in AD. Furthermore, the *trans-* association between SNP rs4420638 and SNRPF had opposite directionality compared to that with APOE levels. These observations suggest that a regulatory mediator is most likely involved. Theoretically, pharmacological targeting of this mediator could cause an increase in SNRPF, which would increase splicing, and potentially decrease amyloid precursor protein levels.

### Pharmacogenetics

A pQTL that harbours a drug target may affect patient response to treatment. For example, we observed a strong, replicated *cis-* association between rs489286 and reduced blood levels of the signalling lymphocytic activation molecule-F7 (SLAMF7) in carriers of the minor allele. This was further confirmed with a SLAMF7-eQTL that showed identical directionality in mRNA sequencing data from lymphoblastoid cell lines (**Supplemental Figure 4**). Furthermore, imputed data identified a strong association between SLAMF7 protein levels and a SNP in the SLAMF7 intron, rs11581248; heterozygous alleles caused severely reduced SLAMF7 levels, and the homozygous minor allele nearly ablated the SLAMF7 protein (Figure 2). SLAMF7 is targeted by the recently FDA-approved cancer drug, Elotuzumab, a humanized monoclonal antibody prescribed for relapsed or refractory multiple myeloma. Previously, a small study on Japanese women indicated an association between rs17313034 (r^2^=0.82 with rs489286) and cervical cancer ^46^; it showed that the minor allele had a protective effect. Taken together, these data suggest the possibility that the response to Elotuzumab treatment may depend on the patient’s rs489286 genotype, and that rs11581248 homozygotes (1.5% of the population) may not respond to Elotuzumab. We cannot exclude the possibility that rs17313034 might affect the SLAMF7 epitope, which could potentially alter SLAMF7-aptamer binding. However, such an alteration might then also affect Elotuzumab binding. This hypothesis can be tested retrospectively in the phase-3 clinical trial cohort ^47^.

### Drug target validation

Plenge *et al*. ^48^ suggested that naturally occurring genetic variance could be used to validate drug targets, based on genotype-phenotype dose-response curves. For example, IL6R was previously proposed as a target for preventing coronary heart disease. Recently, tocilizumab, a humanized antibody that targets IL6R, was developed and is currently approved for treating rheumatoid arthritis. Plenge et al.^48^ required that, for validating drug targets, the gene must include multiple causative variants of known biological function. A large, IL6R Mendelian randomization analysis ^49^ found that SNP rs7529229 was associated with increased IL6R, reduced CRP, reduced fibrinogen, and reduced odds of coronary heart disease (p=1.53×10^-5^). The CardiogramPlus consortium GWAS data confirmed that association (p=1.66×10^-8^). In the present study, we replicated the IL6R-pQTL (rs4129267, r^2^=0.94 with rs7529229). Furthermore, we identified a second SNP in IL6R, rs11804305, which tags an independent and causative variant, since rs4129267 was already established as functional. Therefore, these two SNPs could be used to investigate dose-response curves in the huge dataset available from the IL6R-Mendelian randomization consortium^49^. A similar approach is now feasible for all other protein drug targets that were associated here with multiple pQTLs (for examples see **Supplemental Figure 2**). What is more, the aptamers that were used in this study to target these proteins can readily be used as intermediate readouts in assessing drug responses and in optimizing the efficacy of lead components.

### Application to disease GWAS

Genetic associations with intermediate traits are generally much stronger than associations with disease endpoints, due to their proximity to the causative variant, as shown in our previous GWAS with metabolic traits^50^. Therefore, pQTLs can serve as proxies to fine-map genetic disease associations. This approach can be used to identify potentially causative genes and additional independent genetic signals at a locus identified in a disease-GWAS. In particular, pQTLs can be used to identify true positive associations among associations that do not reach genome-wide significance. For instance, rs12146727 was associated with cardiovascular disease (CVD; p=6.18×10^-5^) in CardiogramPlus and with AD (p=3.10×10^-5^ in the discovery cohort (*st1)*, and p=8.27×10^-6^ in a combined analysis of a larger cohort (*st12comb*)) in the International Genomics of Alzheimer‘s Project. A true positive association requires that the association signal must be strengthened with increasing sample numbers, which was the case for the *st1* and *st12comb* cohorts. In the present study, rs12146727 was replicated as a *cis*-pQTL for complement C1r subcomponent (C1R), and it was replicated as a *trans*-pQTL for complement C1q subcomponent (C1QA/C1QB/C1QC). These associations support growing evidence that suggests that the complement system plays a role in both AD and CVD ^51^. This example also shows how variants that associate with a disease endpoint could generate new hypotheses about the role of the co-associated protein(s) in the disease aetiology. Note that even when these variants have small effect sizes or odds ratios, the related pathways may show large responses to pharmaceutical alteration of these proteins. Hypotheses generated with this approach can be directly tested in animal models, and the existing aptamers can be potentially used as intermediate functional readouts in the drug development process.

## DISCUSSION

Genetic studies with yeast^52^ and lymphoblastoid cell lines^53–55^ have indicated that cellular protein levels are under strong genetic control. This control was confirmed by the discovery of many *cis-* acting genetic variants in the human blood proteome^4–9^. Here, we described the first large-scale proteomics GWAS on blood plasma proteins derived from a human population. This GWAS represented over 1.1 million individual aptamer binding experiments. By design, our panel of more than 1100 aptamer targets was highly enriched in biomedically-relevant blood circulating proteins, which was reflected in the large overlap of pQTLs with risk loci identified in disease-GWAS. The data generated from this study can be used in future investigations to identify and validate causative variants identified in disease GWAS. For instance, aptamer-based read-outs can be used as intermediate traits in CrispR-based experiments, as exemplified with the FTO-obesity association described by Claussnitzer et al.^56,57^.

Surprisingly, we found that up to 60% of the naturally occurring variance in in the blood plasma levels of essential proteins could be explained by two or more independent variants of a single gene located on another chromosome. We identified strong genetic associations with intermediate, and most likely functional, traits related to proteins involved in pathways to complex disorders. While many of our associations connect these proteins to disease pathway through shared association, it should be borne in mind that confounding is always a possibility that needs to be ruled out by further experimentation. These findings can be used in future studies to establish dose-response curves for drug-target validation^48^. A greater understanding of the genetic control of circulating levels of protein drug targets and biomarkers may improve pharmaceutical interventions and clinical trials. Because all the aptamers used in this study were synthetically generated and well defined, they can be readily developed into specific assays for precise clinical applications. For instance, the SLAMF7-binding aptamer identified here, which revealed a potential genetic effect on Elotuzumab, may be developed directly into a clinical assay for identifying differential responders to immunotherapy. Moreover, this concept can be generalized to other drug targets and biomarkers.

Despite the variable baseline conditions among the participants of the replication cohort (not fasting, high prevalence of diabetes, and multiple ethnic backgrounds), we replicated 82% of all sufficiently powered associations. This result emphasized the robustness of the replicated associations. It also suggests that many of the Bonferroni-significant associations that could not be replicated in our QMDiab cohort may be replicated in future studies. In this study, about 20% of all assayed proteins had a significant pQTL association. This number is expected to increase in future more highly powered studies.

Genetic variance in a protein sequence may affect its higher order structure, and thus, its aptamer-binding affinity. Similarly, alterations in protein structure may affect binding and specificity in immunoassay-based methods and protein mass in targeted MS-based methods. These issues remain to be addressed. In this study, we showed that structural-based epitope effects on a particular pQTL may be identified or ruled out in several ways, including: allele-specific transcription analysis (e.g., CPNE1, **Supplemental Figure 5**); co-associated eQTLs, particularly when they include multiple genetic variants (e.g., ERAP1, Figure 4); the absence of correlated SNPs that alter the protein structure (e.g., based on SNiPA-annotated 1000 Genomes data), or replication on different platforms (**Supplemental Table 2-5**). We did not directly follow up on our results using mass spectroscopy. However, Ngo et al. ^58^ recently showed that response curves for selected aptamer-enriched proteins were linear over a wide dynamic range of spiked protein concentrations. Most importantly, *trans-* associations, which were a central focus of this study, are not affected by this type of potential artefact.

In summary, our GWAS demonstrated the power of linking the genome to disease endpoints via the blood proteome. As we have shown at the examples, mining of our data can reveal a plethora of new insights into biological processes and provide a wealth of functional information that is beyond the focus of a single publication. Therefore, we have provided additional interpretations of selected loci in **Supplemental Note 1.** Furthermore, our complete results are freely accessible for further analyses of pQTL associations with past and future disease GWAS, on an integrated web-server at http://proteomics.gwas.eu.

## METHODS

### Study population

<u>KORA</u>: The KORA F4 study is a population-based cohort of 3,080 subjects living in southern Germany. Study participants were recruited between 2006 and 2008 comprising individuals who, at that time, were aged 32–81 years. KORA F4 is the follow-up study of the KORA S4 survey conducted in 1999–2001 (4,261 participants). The study design, standardized sampling methods and data collection (medical history, questionnaires, anthropometric measurements) have been described in detail elsewhere (^11^ and references therein). For this study, 1,000 individuals were randomly selected from a subset of 1,800 already deeply phenotyped KORA F4 study participants ^1,59,60^. All study participants gave written informed consent and the study was approved by the Ethics Committee of the Bavarian Medical Association. <u>QMDiab</u>: The Qatar Metabolomics Study on Diabetes (QMDiab) is a cross-sectional case-control study that was conducted between February and June 2012 at the Dermatology Department of HMC in Doha, Qatar. QMDiab has been described previously and comprises male and female participants in near equal proportions, aged between 23 and 71 years, mainly from Arab, South Asian and Filipino descent ^12^. The initial study was approved by the Institutional Review Boards of HMC and Weill Cornell Medicine – Qatar (WCM-Q) (research protocol #11131/11). Written informed consent was obtained from all participants.

### Blood sampling

<u>KORA</u>: Blood samples for omics-analyses and DNA extraction were collected between 2006 and 2008 as part of the KORA F4 follow-up. To avoid variation due to circadian rhythm, blood was drawn in the morning between 08:00 and 10:30 after a period of at least 10 h overnight fasting. Blood was collected without stasis and was kept at 4 °C until centrifugation. The material was then centrifuged for 10 min (2,750g at 15 °C). Plasma samples were aliquoted and stored at −80 °C until assayed on the SOMAscan platform. <u>QMDiab</u>: Non-fasting plasma specimens were collected in the afternoon, after the general operating hours of the morning clinic, and processed using standardized protocols. Cases and controls were collected as they became available, in a random pattern and at the same location using identical protocols, instruments and study personnel. Samples from cases and controls were processed in the lab blinded to their identity. After collection the samples were stored on ice for transportation to WCM-Q. Within six hours after sample collection all samples were centrifuged at 2,500g for 10 minutes, aliquoted, and stored at -80°C until analysis.

### Genotyping

<u>KORA</u>: The Affymetrix Axiom Array was used to genotype 3,788 participants of the KORA S4 study. After thorough quality control (total genotyping rate in the remaining SNPs was 99.8%) and filtering for minor allele frequency >1%, a total of 509,946 autosomal SNPs was kept for the GWAS analysis. 1000 Genomes Project-imputed genotypes were used for fine mapping and generation of regional association plots (**Supplemental Figure 6**). KORA genotype data has been reported previously with many GWAS studies and was used here as-provided by the consortium. We therefore do not repeat details here ^11^. <u>QMDiab</u>: DNA was extracted from 359 samples from QMDiab and genotyped by the WCM-Q genomics core facility using the Illumina Omni 2.5 array (version 8). High quality genotype data of 2,338,671 variants was obtained for 353 samples; data for 6 samples was excluded due to a low overall call rate (<90%). Removal of duplicate variants left 2,327,362 variants. 134,830 variants were removed due to missing genotype data (PLINK option --geno 0.02), leaving 2,192,532 variants. 941,058 variants were removed due to minor allele threshold (PLINK option --maf 0.05), leaving 1,251,474 variants. 28,175 variants were removed due to violation of Hardy-Weinberg equilibrium (PLINK option -- hwe 1E-6), leaving 1,223,299 variants. Of these variants, 1,221,345 were autosomal variants. The total genotyping rate of these remaining variants was 99.7%.

### Proteomics measurements

<u>KORA</u>: The SOMAscan platform was used to quantify protein levels. It has been described in detail before ^5,10,61–65^. Briefly, undepleted EDTA-plasma is diluted into three dilution bins (0.05%, 1%, 40%) and incubated with bin-specific collections of bead-coupled SOMAmers in a 96-well plate format. Subsequent to washing steps, bead-bound proteins are biotinylated and complexes comprising biotinylated target proteins and fluorescence-labeled SOMAmers are photocleaved off the bead support and pooled. Following recapture on streptavidin beads and further washing steps, SOMAmers are eluted and quantified as a proxy to protein concentration by hybridization to custom arrays of SOMAmer-complementary oligonucleotides. Based on standard samples included on each plate, the resulting raw intensities are processed using a data analysis work flow including hybridization normalization, median signal normalization and signal calibration to control for inter-plate differences. 1,000 blood samples from the KORA F4 study were sent to SomaLogic Inc. (Boulder Colorado, USA) for analysis. Two of the shipped samples were incorrectly pulled from the bio-bank and had no corresponding genotype data, one sample failed SOMAscan QC, leaving a total of 997 samples from 483 males and 514 females for analysis. Data for 1,129 SOMAmer probes (SOMAscan assay V3.2) was obtained for these samples. Five of the probes failed SOMAscan QC, leaving a total of 1,124 probes for analysis (**Supplemental Table 1**). <u>QMDiab</u>: 352 samples from QMDiab were analyzed at the WCM-Q proteomics core, 338 of which overlapped with samples that were also genotyped. No samples were excluded. Protocols and instrumentation were provided and certified using reference samples by SomaLogic Inc.. Experiments were conducted under supervision of SomaLogic personnel. No samples or probe data was excluded.

### GWAS discovery study

We used PLINK (version 1.90b3w, Shaun Purcell, Christopher Chang, https://www.cog-genomics.org/plink2) ^66^ to fit linear models to inverse-normalized probe levels, using age, gender, and body mass index (BMI) as covariates. R version 3.1.3 (R: A language and environment for statistical computing, R Foundation for Statistical Computing, Vienna, Austria, 2015, www.R–project.org/) was used for data organization, plotting and additional statistical analyses outside of the actual GWAS, including computation of linear regression models based on raw and log-scaled data. We retained all SNP-probe associations with p-values < 10^-5^ (14,647 associations covering 3,169 unique SNPs and 1,118 unique probes, **Supplemental Data 2**, see **Supplemental Figure 7** for a Manhattan plot with all associations). Genomic inflation was low (mean=1.0035, max=1.0251). Therefore no correction for genomic control was applied. To define genetically independent association loci, we lumped in a first step for every given probe all associations with correlated SNPs (LD r^2^>0.1, window size 10Mb) and retrieved the SNP that had the strongest association at the sentinel for this probe association. We then grouped all highly correlated sentinel SNPs (LD r^2^>0.9, window size 10Mb) into single loci. We report uncorrected association p-values throughout this paper. In all statistical analyses we require a nominal significance level of 0.05 (alpha error 5%). Multiple hypotheses testing is accounted for by using conservative Bonferroni correction of the significance level, based on the number of tested SNPs (509,946) and probes (1,124), resulting in a genome- and proteome-wide significance level of p<8.72×10^-11^ (0.05/509,946/1,124). 451 loci had at least one Bonferroni significant sentinel association. For each of these 451 loci, we also considered all probe associations with that same SNP at a significance level of p<9.86×10^-8^ (0.05/451/1,124) as Bonferroni significant. This resulted in a total of 539 genetic association signals (284 unique probes), each represented by a sentinel SNP and one or more sentinel probes, including 391 *cis-* associations (defined by a SNP closer than 10Mb from the gene boundaries, 202 unique probes, 18.5% of all 1,090 autosomal probes) and 148 *trans-* associations (96 unique probes, 8.5% of all 1,124 probes). Box plots and association statistics using alternative data scaling methods (raw, log-normal, inverse-normal) are provided as **Supplemental Figure 8**. Annotations of the genetic loci using the SNiPA web-server are provided as **Supplemental Figure 9**. Regional association plots are in **Supplemental Figure 6**.

### Replication

Using the QMDiab proteomics data as dependent variables, linear regression models were fitted with PLINK (version 1.90, version b3w), using age, sex, BMI, diabetes state, the first three principal components (PCs) of the genotype data and the first three PCs of the proteomics data as covariates. The first three genetic PCs separate the three major ethnicities and the three proteomics PCs account for variability introduced by a low degree of cell haemolysis. Genomic inflation was low (mean=1.020, max=1.116). Therefore no correction for genomic inflation was applied. A tag SNP for replication of each of the 451 Bonferroni significant loci was selected by using the strongest association among all imputed SNPs in the discovery study which had a correlation of r^2^>0.8 with the sentinel SNP. 387 tag SNPs for the 451 originally identified SNPs could be identified for replication, covering 462 SNP-probe pairs out of the originally identified 539 SNP-probe pairs. To consider an association as replicated, we therefore required p<1.08×10^-4^ (0.05/462). 234 (50.6%) of the 462 attempted replications fully replicated at a Bonferroni level of significance (p<0.05/462), 384 (83.1%) displayed nominal significance (p<0.05). Out of 84 non replicated associations that were nominally significant and that had a MAF<30% in both cohorts, only one association displayed a discordant trend. We estimated the statistical power for the replication by sampling: For each association we randomly selected without replacement 338 individuals from the KORA cohort and computed the p-value of association on that subset. We repeated this 100 times and report the 5^th^ largest p-value from this empirical distribution as the p-value that can be expected to be obtained with 95% power (p95). Based on this analysis we found that 208 SNP-probe pairs with a suitable tag SNP in QMDiab had 95% replication power (p95<1.08×10^-4^). 171 of these 208 (82.2%) fully replicated in QMDiab, 198 (95.2%) displayed at least nominal significance.

### Replication of previous *cis-* associations with SOMAscan technology

Using the SOMAscan platform on 100 samples, Lourdusamy et al. ^5^ reported 60 *cis-* associations at a false discovery rate (FDR) of 5%. 48 of these associations had a suitable tag SNP in the genotyped KORA data set (r^2^>0.5) or association data on an imputed SNP that allowed for replication. 34 of these SNPs were replicated (p<0.05/120, conservatively accounting for replication attempts on tagged and imputed SNPs). The first non-replicated association had rank 21 (**Supplemental Table 3**).

### Replication of immunoassay based *cis-* associations

Kim et al. ^7^ reported 28 *cis-* associations for 27 analytes in the ADNI cohort using plasma proteomic data by multiplex immunoassay on the Myriad Rules Based Medicine (RBM) Human DiscoveryMAP panel v1.0 on the Lumine×100 platform. Out of 17 associations that had overlapping probes, thirteen were replicated (p<0.05/17) (**Supplemental Table 4**). Melzer et al. ^4^ tested 40 protein levels determined by immunoassay and reported 8 pQTLs. We replicated their second strongest association with IL6R. Their strongest association is at the ABO locus with TNFa. However, TNFa is not on the SOMAscan panel. We also found numerous other strong associations at the ABO locus. Two weak associations of Melzer et al. (SHBG and CRP) were not significant in our study. The remaining associations from Melzer et al. were not targeted by our panel. Enroth et al. ^9^ used a multiplexed immunoassay and reported 23 *cis-* pQTLs. Two of the proteins (four associations) targeted by their assay were not covered by our panel (VEGF-D, Ep-CAM) and one was located on the X-chromosome and not tested for *cis-* association here (CD40-L). Of the remaining 18 *cis-* pQTLs, twelve associations were replicated (p<0.05/18) (**Supplemental Table 5**).

### Replication of MS-based *cis-* associations

Johansson^6^ reported five associations and we found *cis-* pQTLs for four of them (Haptoglobin (HP), Alpha-1-antitrypsin (SERPINA1), Alpha-2-HS-glycoprotein (AHSG), and APOE isoform 2). We did not, however, find a *cis-* pQTL for Complement C3 (C3), despite the fact that several isoforms were targeted by our panel. Liu et al. ^8^ reported 13 statistically significant association of MS-derived protein levels. Four of their proteins (FCN2, ITIH4, KNG1, PON1) were targeted by our panel, and for two of them we found a *cis-* pQTL in our study (KNG1, FCN2). Wu et al. ^53^ reported protein associations using MS in lymphoblastoid cell lines. However, this study was targeting intracellular proteins and had no overlap with blood circulating proteins targeted by our panel and could therefore not be used for replication.

### Proteome annotation

We used SOMAmer probe identifiers as primary protein identifiers (**Supplemental Table 1**). Some SOMAmer targets map to multiple Uniprot identifiers (N=41). They either refer to protein complexes (N=32) that are encoded at multiple genome loci, or to different variants of a same protein encoded at a single gene locus (N=9). 1,090 SOMAmer targets were encoded on autosomal chromosomes, 30 targets were encoded on the X-chromosome, and four targeted viral proteins. Genome positions for all SOMAmer targets were retrieved from Ensembl (http://www.ensembl.org) using probe-specific Uniprot identifiers provided by SomaLogic. We used Ingenuity Pathway Analysis (IPA) (http://www.ingenuity.com/products/ipa) to retrieve additional information related to each probe. IPA provides a rich expert-curated knowledgebase of literature based protein-related information and requires unique mapping to protein identifiers. Forty-one probes were present that target proteins with multiple Uniprot IDs, and these were excluded from the IPA analysis. A further 29 probes were excluded because multiple probes target a same protein. Eight Uniprot IDs could not be mapped by IPA (LAG-1, LD78-beta, NKG2D, HSP 70, and four viral proteins). Ultimately, 1,045 probes with unique Uniprot IDs remained for the IPA annotation.

### Functional annotation of the associations

We used the SNiPA server (v3.1, http://snipa.org) to annotate 435 out of our 451 lead SNPs (**Supplemental Figure 9**). 16 SNPs were not available in the SNiPA database. SNiPA provides annotations for all SNPs that are in linkage disequilibrium (R^2^>0.8) with and no more than 500kb distant from a sentinel SNP using genome assembly data based on GRCh37.p13, Ensembl version 82, 1000-Genomes (phase 3, version 5) data, and GTEx (release 4) eQTL associations for 13 tissues (see columns SNIPA_… in **Supplemental Data 2**). SNiPA also provides primary effect predictions using the Ensembl VEP tool ^67^ and all GWAS association data from the GWAS catalog ^68^, metabotype associations (http://gwas.eu, and dbGaP(columnsGWAS_TRAITS, mGWAS_TRAITS, dbGaP_TRAITS, accessed October 2015). Updates can be retrieved online at http://snipa.org using the block-annotation tool.

### Methylation GWAS

Data was downloaded from http://genenetwork.nl/biosqtlbrowser (accessed 9 Feb. 2016). This web server accompanies a manuscripts entitled ‘Disease variants alter transcription factor levels and methylation of their binding sites‘, by Bonder & Luijk *et al*., and ‘Unbiased identification of regulatory modifiers of genetic risk factors‘, by Zhernakova *et al*. Both papers have been submitted to Nature Genetics. The manuscript was accessed on bioRxiv as a preprint, first posted online November 30, 2015; doi: http://dx.doi.org/10.1101/033084. Tagging SNPs (r^2^>0.8) were identified using SNP-SNP correlations from KORA imputed genotypes and meQTLs with p-values < 10^−9^ were retrieved.

### Metabolomics GWAS

GWAS associations with metabolic traits (mQTLs) were obtained using the GWAS-server (http://gwas.eu). This web-server provides access to raw association data from the studies by Suhre et al. ^1^, Shin et al. ^20^, and Raffler et al. ^69^. Extracted associations were limited to p-value < 10^-8^ and further requiring p-gain > 10^4^ for ratios ^44^ (**Supplemental Table 6**).

### Disease GWAS lookup

We used SNiPA to identify overlapping disease-GWAS entries. We further downloaded publically available association data for 70 clinically relevant traits from the web-sites of 14 large disease GWAS consortia, based on a list of available GWAS, and downloaded from https://www.med.unc.edu/pgc/downloads. **Supplemental Data 4** provides a list with links to all downloaded files and the corresponding publications.

### QMDiab total plasma N-glycosylation

Unthawed aliquots of identical samples as for the proteome analysis were sent to Genos Ltd. (Zagreb, Croatia) for analysis using ultra-performance liquid chromatography and liquid chromatography mass spectrometry glycoprofiling as follows: <u>Total plasma N-glycan release and labeling.</u> Glycans were released from total plasma proteins and labeled as described previously ^70^. Briefly, 10 μL plasma sample was denatured with the addition of 20 μl 2% (w/v) SDS (Invitrogen, USA) and N-glycans were released with the addition of 1.2 U of PNGase F (Promega, USA). The released N-glycans were labeled with 2-aminobenzamide (Sigma-Aldrich, USA). Free label and reducing agent were removed from the samples using hydrophilic interaction liquid chromatography solid-phase extraction. 0.2 μm 96-well GHP filter-plate (Pall Corporation, USA) was used as stationary phase. Samples were loaded into the wells and after a short incubation washed 5x with cold 90% ACN. Glycans were eluted with 2 × 90 μL of ultrapure water after 15 min shaking at room temperature, and combined eluates were stored at −20°C until use. <u>Total plasma N-glycome UPLC analysis</u>. Total plasma N-glycans were analyzed by hydrophilic interaction ultra-performance liquid chromatography (HILIC-UPLC) as described previously ^70^. Briefly, fluorescently labeled N-glycans were separated on an Acquity UPLC instrument (Waters, USA) using excitation and emission wavelengths of 250 and 428 nm, respectively. Labeled N-glycans were separated on a Waters BEH Glycan chromatography column, 150 × 2.1 mm i.d., 1.7 μm BEH particles, with 100 mM ammonium formate, pH 4.4, as solvent A and acetonitrile (ACN) (Fluka, USA) as solvent B. The separation method used a linear gradient of 30-47% solvent A at flow rate of 0.56 mL/min in a 23 min analytical run (**Supplemental Figure 10**).

### mRNA sequencing and allele specific transcription analysis

Lappalainen et al. ^71^ report mRNA sequencing of 462 lymphoblastoid cell lines of the 1000 Genomes Project. RNA sequencing data was downloaded from EBI (http://www.ebi.ac.uk/Tools/geuvadis-das). We used CLCBio genomics workbench (Qiagen Inc.) to align reads, calculate RKPM (Reads Per Kilobase of transcript per Million mapped reads) values and analyse the data.

### Gaussian Graphical Network (GGM) construction

Using raw (unscaled) data, regressing out age + gender + bmi (997 samples with data for 1,124 variables) we computed a GGM using the ggm.estimate.pcor function from the R GeneNet package. The estimated optimal shrinkage intensity lambda (correlation matrix) was 0.187. We obtained 3,943 GGM edges connecting 1,092 protein nodes with Bonferroni significant partial correlation coefficients (p< 7.9×10^-8^ = 0.05/(1,124*1,123/2)), provided as **Supplemental Data 3**. We added SNP-probe association edges connecting 451 genetic loci to 539 proteins. We further added SNP-disease association edges using all associations reported in the GWAS catalogue (identified using SNiPA at LD r^2^>0.8). We also added all SNP-disease associations from 84 clinically relevant traits of 14 large GWAS consortia that had a p-value P<10^-8^. These SNP-disease edges were further manually curated to ascertain unique SNP-disease pairs. SNP association to disease-related protein levels were excluded.

### Data Availability

All summary statistics and association data are freely available, accessible online on an integrated web-server at http://proteomics.gwas.eu. A fully functional version of the web-server can also be freely downloaded from this link for local installation and network-free usage (HTML5-based, no extra software is required). The informed consent given by the study participants does not cover posting of participant level phenotype and genotype data in public databases. However, data are available upon request from KORA-gen (http://epi.helmholtz-muenchen.de/kora-gen). Requests are submitted online and are subject to approval by the KORA board.

## Acknowledgements

This work was supported by ‘Biomedical Research Program’ funds at Weill Cornell Medicine in Qatar, a program funded by the Qatar Foundation. The statements made herein are solely the responsibility of the authors. M. Arnold was supported by the Helmholtz cross-program topic “Metabolic Dysfunction”. D. Mook-Kanamori was supported by Dutch Science Organization (ZonMW-VENI Grant 916.14.023). The KORA study was initiated and financed by the Helmholtz Zentrum München – German Research Center for Environmental Health, which is funded by the German Federal Ministry of Education and Research (BMBF) and by the State of Bavaria. Furthermore, KORA research was supported within the Munich Center of Health Sciences (MC-Health), Ludwig-Maximilians-Universität, as part of LMUinnovativ. The KORA-Study Group consists of A. Peters (speaker), J. Heinrich, R. Holle, R. Leidl, C. Meisinger, K. Strauch, and their co-workers, who are responsible for the design and conduct of the KORA studies. We gratefully acknowledge the contribution of all members of field staff conducting the KORA F4 study. We thank the staff from HMC dermatology department and WCM-Q clinical research core for their contribution to QMDiab. We thank Brian Sellers from SomaLogic for support with measuring the QMDiab samples at WCM-Q. We acknowledge free access to summary statistics provided by the GWAS consortia listed in **Supplemental Data 4**. Most of all, we thank all study participants of KORA and QMDiab for their invaluable contributions to this study.

### Author contributions

Jointly supervised research: KS, JG, CG

Conceived and designed the experiments: KS, JG, CG

Performed the experiments: RE, JG, EKA, YAM, JM, HS, GL, MP,

Performed statistical analysis: KS, CG, AL

Analysed the data: KS, AB, RJC, JG, GT, MA, CG, GK, AL, JR

Contributed reagents/materials/analysis tools: KS, MAS, MA, HG, GK, AP, JR, KStr, DOM, RKD, LG

Wrote the paper: KS

All authors discussed the results and reviewed the final manuscript.

MA, AB, RJC, RE, AL, JR, HS, and GT contributed equally to this work and are listed in alphabetic order.

## REFERENCES

1. Suhre, K. et al. Human metabolic individuality in biomedical and pharmaceutical research. Nature 477, 54–60 (2011).

2. Stunnenberg, H. G. & Hubner, N. C. Genomics meets proteomics: Identifying the culprits in disease. Hum. Genet. 133, 689–700 (2014).

3. Gutierrez-Arcelus, M., Rich, S. S. & Raychaudhuri, S. Autoimmune diseases — connecting risk alleles with molecular traits of the immune system. Nat. Rev. Genet. 17, 160–174 (2016).

4. Melzer, D. et al. A genome-wide association study identifies protein quantitative trait loci (pQTLs). PLoS Genet. 4, e1000072 (2008).

5. Lourdusamy, A. et al. Identification of cis-regulatory variation influencing protein abundance levels in human plasma. Hum. Mol. Genet. 21, 3719–3726 (2012).

6. Johansson, Å. et al. Identification of genetic variants influencing the human plasma proteome. Proc. Natl. Acad. Sci. U. S. A. 110, 4673–8 (2013).

7. Kim, S. et al. Influence of Genetic Variation on Plasma Protein Levels in Older Adults Using a Multi-Analyte Panel. PLoS One 8, (2013).

8. Liu, Y. et al. Quantitative variability of 342 plasma proteins in a human twin population. Mol. Syst. Biol. 11, 786 (2015).

9. Enroth, S., Johansson, A., Enroth, S. B. & Gyllensten, U. Strong effects of genetic and lifestyle factors on biomarker variation and use of personalized cutoffs. Nat. Commun. 5, 4684 (2014).

10. Gold, L. iet al. Aptamer-based multiplexed proteomic technology for biomarker discovery. PLoS One 5, (2010).

11. Wichmann, H.-E., Gieger, C. & Illig, T. KORA-gen--resource for population genetics, controls and a broad spectrum of disease phenotypes. Gesundheitswesen 67 Suppl 1, S26–30 (2005).

12. Mook-Kanamori, D. O. et al. 1,5-Anhydroglucitol in saliva is a noninvasive marker of short-term glycemic control. J. Clin. Endocrinol. Metab. 99, (2014).

13. Arnold, M., Raffler, J., Pfeufer, A., Suhre, K. & Kastenmüller, G. SNiPA: an interactive, genetic variant-centered annotation browser.

14. Dichgans, M. et al. Shared genetic susceptibility to ischemic stroke and coronary artery disease : A genome-wide analysis of common variants. Stroke 45, 24–36 (2014).

15. Qi, L. et al. Genetic variants in ABO blood group region, plasma soluble E-selectin levels and risk of type 2 diabetes. Hum. Mol. Genet. 19, 1856–1862 (2010).

16. Dentali, F. et al. Non-O blood type is the commonest genetic risk factor for VTE: Results from a meta-analysis of the literature. Semin. Thromb. Hemost. 38, 535–547 (2012).

17. Wolpin, B. M. et al. Pancreatic cancer risk and ABO blood group alleles: Results from the Pancreatic Cancer Cohort Consortium. Cancer Res. 70, 1015–1023 (2010).

18. Fry, A. E. et al. Common variation in the ABO glycosyltransferase is associated with susceptibility to severe Plasmodium falciparum malaria. Hum. Mol. Genet. 17, 567–76 (2008).

19. Koyama, H. et al. Platelet P-selectin expression is associated with atherosclerotic wall thickness in carotid artery in humans. Circulation 108, 524–529 (2003).

20. Shin, S.-Y. et al. An atlas of genetic influences on human blood metabolites. Nat. Genet. 46, 543–50 (2014).

21. Altmaier, E. et al. Metabolomics approach reveals effects of antihypertensives and lipid-lowering drugs on the human metabolism. Eur. J. Epidemiol. 29, 325–336 (2014).

22. Chan, B., Yuan, H. T., Ananth Karumanchi, S. & Sukhatme, V. P. Receptor tyrosine kinase Tie-1 overexpression in endothelial cells upregulates adhesion molecules. Biochem. Biophys. Res. Commun. 371, 475–479 (2008).

23. Morrow, D., Cullen, J. P., Cahill, P. A. & Redmond, E. M. Cyclic strain regulates the Notch/CBF-1 signaling pathway in endothelial cells: Role in angiogenic activity. Arterioscler. Thromb. Vasc. Biol. 27, 1289–1296 (2007).

24. Zanetti, A. et al. Vascular endothelial growth factor induces Shc association with vascular endothelial cadherin: A potential feedback mechanism to control vascular endothelial growth factor receptor-2 signaling. Arterioscler. Thromb. Vasc. Biol. 22, 617–622 (2002).

25. Nilsson, I. et al. VEGF receptor 2/-3 heterodimers detected in situ by proximity ligation on angiogenic sprouts. EMBO J. 29, 1377–1388 (2010).

26. Lu, K. V. et al. VEGF Inhibits Tumor Cell Invasion and Mesenchymal Transition through a MET/VEGFR2 Complex. Cancer Cell 22, 21–35 (2012).

27. Stannard, A. K. et al. Vascular endothelial growth factor synergistically enhances induction of E-selectin by tumor necrosis factor-α. Arterioscler. Thromb. Vasc. Biol. 27, 494–502 (2007).

28. Smith, G. a. et al. Vascular endothelial growth factors: multitasking functionality in metabolismhealth and disease. J. Inherit. Metab. Dis. 753–763 (2015). doi:10.1007/s10545-015-9838-4

29. He, M. et al. A genome wide association study of genetic loci that influence tumour biomarkers cancer antigen 19-9, carcinoembryonic antigen and alpha fetoprotein and their associations with cancer risk. Gut 63, 143–51 (2013).

30. Yue, T. et al. Identification of blood-protein carriers of the CA 19-9 antigen and characterization of prevalence in pancreatic diseases. Proteomics 11, 3665–3674 (2011).

31. Narimatsu, H. et al. Lewis and Secretor Gene Dosages Affect CA19-9 and DU-PAN-2 Serum Levels in Normal Individuals and Colorectal Cancer Patients Lewis and Secretor Gene Dosages Affect CA19-9 and DU-PAN-2 Serum Levels in Normal Individuals and Colorectal Cancer Patients1. Cancer Res. 58, 512–518 (1998).

32. Vestergaard, E. M. et al. Reference values and biological variation for tumor marker CA 19-9 in serum for different Lewis and secretor genotypes and evaluation of secretor and Lewis genotyping in a Caucasian population. Clin. Chem. 45, 54–61 (1999).

33. Ritchie, G. E. et al. Glycosylation and the complement system. Chem. Rev. 102, 305–319 (2002).

34. Arnold, J. N. et al. Interaction of mannan binding lectin with ??2 macroglobulin via exposed oligomannose glycans: A conserved feature of the thiol ester protein family? J. Biol. Chem. 281, 6955–6963 (2006).

35. Song, G. G. et al. Meta-analysis of functional MBL polymorphisms. Z. Rheumatol. 73, 657–664 (2014).

36. Stahl, E. A. et al. Bayesian inference analyses of the polygenic architecture of rheumatoid arthritis. Nat. Genet. 44, 483–9 (2012).

37. Evans, D. M. et al. Interaction between ERAP1 and HLA-B27 in ankylosing spondylitis implicates peptide handling in the mechanism for HLA-B27 in disease susceptibility. Nat. Genet. 43, 761–767 (2011).

38. Zervoudi, E. et al. Rationally designed inhibitor targeting antigen-trimming aminopeptidases enhances antigen presentation and cytotoxic T-cell responses. Proc. Natl. Acad. Sci. U. S. A. 110, 19890–19895 (2013).

39. Hattori, A. & Tsujimoto, M. Endoplasmic reticulum aminopeptidases: Biochemistry, physiology and pathology. J. Biochem. 154, 219–228 (2013).

40. Martín-Esteban, A., Gomez-Molina, P., Sanz-Bravo, A. & De Castro, J. A. L. Combined effects of ankylosing spondylitis-associated erap1 polymorphisms outside the catalytic and peptide-binding sites on the processing of natural HLA-B27 ligands. J. Biol. Chem. 289, 3978–3990 (2014).

41. Reeves, E., Edwards, C. J., Elliott, T. & James, E. Naturally Occurring ERAP1 Haplotypes Encode Functionally Distinct Alleles with Fine Substrate Specificity. J Immunol 191, 35–43 (2013).

42. Hellwage, J. et al. Functional properties of complement factor H-related proteins FHR-3 and FHR-4: Binding to the C3d region of C3b and differential regulation by heparin. FEBS Lett. 462, 345– 352 (1999).

43. Frimat, M. et al. Complement activation by heme as a secondary hit for atypical hemolytic uremic syndrome Complement activation by heme as a secondary hit for atypical hemolytic uremic syndrome. 122, 282–292 (2013).

44. Petersen, A.-K. et al. On the hypothesis-free testing of metabolite ratios in genome-wide and metabolome-wide association studies. BMC Bioinformatics 13, 120 (2012).

45. Bai, B. et al. U1 small nuclear ribonucleoprotein complex and RNA splicing alterations in Alzheimer’s disease. Proc. Natl. Acad. Sci. U. S. A. 110, 16562–7 (2013).

46. Kiyonor Miura, Hiroyuki Mishima, Akira Kinoshita, Chisa Hayashida, Shuhei Abe, Katsushi Tokunaga, Hideaki Masuzaki, K. Y. Genome-wide association study of HPV-associated cervical cancer in Japanese women. J. Med. Virol. 1158, 1153–1158 (2014).

47. Lonial, S. et al. Elotuzumab Therapy for Relapsed or Refractory Multiple Myeloma. N. Engl. J. Med. 373, 621–31 (2015).

48. Plenge, R. M., Scolnick, E. M. & Altshuler, D. Validating therapeutic targets through human genetics. Nat. Rev. Drug Discov. 12, 581–594 (2013).

49. Interleukin-6 Receptor Mendelian Randomisation Analysis (IL6R MR) Consortium. The interleukin-6 receptor as a target for prevention of coronary heart disease: a mendelian randomisation analysis. Lancet 379, 1214–1224 (2012).

50. Suhre, K. & Gieger, C. Genetic variation in metabolic phenotypes: study designs and applications. Nature Reviews Genetics 13, 759–769 (2012).

51. Shen, Y., Yang, L. & Li, R. What does complement do in Alzheimer’s disease? Old molecules with new insights. Transl. Neurodegener. 2, 21 (2013).

52. Foss, E. J. et al. Genetic basis of proteome variation in yeast. Nat. Genet. 39, 1369–1375 (2007).

53. Wu, L. et al. Variation and genetic control of protein abundance in humans. Nature 499, 79–82 (2013).

54. Hause, R. J. et al. Identification and Validation of Genetic Variants that Influence Transcription Factor and Cell Signaling Protein Levels. Am. J. Hum. Genet. 95, 194–208 (2014).

55. Garge, N. et al. Identification of Quantitative Trait Loci Underlying Proteome Variation in Human Lymphoblastoid Cells. Mol. Cell. Proteomics 9, 1383–1399 (2010).

56. Claussnitzer, M. et al. Leveraging cross-species transcription factor binding site patterns: From diabetes risk loci to disease mechanisms. Cell 156, 343–358 (2014).

57. Claussnitzer, M. et al. FTO Obesity Variant Circuitry and Adipocyte Browning in Humans. N. Engl. J. Med. 373, 895–907 (2015).

58. Ngo, D. et al. Aptamer-Based Proteomic Profiling Reveals Novel Candidate Biomarkers and Pathways in Cardiovascular Disease. Circulation 134, 270–285 (2016).

59. Illig, T. et al. A genome-wide perspective of genetic variation in human metabolism. Nat. Genet. 42, 137–141 (2010).

60. Petersen, A.-K. K. et al. Epigenetics meets metabolomics: an epigenome-wide association study with blood serum metabolic traits. Hum. Mol. Genet. 23, 534–545 (2014).

61. Kraemer, S. et al. From SOMAmer-based biomarker discovery to diagnostic and clinical applications: A SOMAmer-based, streamlined multiplex proteomic assay. PLoS One 6, (2011).

62. Hathout, Y. et al. Large-scale serum protein biomarker discovery in Duchenne muscular dystrophy. Proc. Natl. Acad. Sci. 112, 201507719 (2015).

63. Sattlecker, M. et al. Alzheimer’s disease biomarker discovery using SOMAscan multiplexed protein technology. Alzheimers. Dement. 10, 724–34 (2014).

64. Kiddle, S. J. et al. Candidate blood proteome markers of Alzheimer’s disease onset and progression: a systematic review and replication study. J. Alzheimers. Dis. 38, 515–31 (2014).

65. Menni, C. et al. Circulating Proteomic Signatures of Chronological Age. Journals Gerontol. Ser. A Biol. Sci. Med. Sci. 1–9 (2014). doi:10.1093/gerona/glu121

66. Chang, C. C. et al. Second-generation PLINK: rising to the challenge of larger and richer datasets. Gigascience 4, 7 (2015).

67. McLaren, W. et al. Deriving the consequences of genomic variants with the Ensembl API and SNP Effect Predictor. Bioinformatics 26, 2069–2070 (2010).

68. Hindorff, L. A. et al. Potential etiologic and functional implications of genome-wide association loci for human diseases and traits. Proc. Natl. Acad. Sci. U. S. A. 106, 9362–7 (2009).

69. Raffler, J. et al. Genome-Wide Association Study with Targeted and Non-targeted NMR Metabolomics Identifies 15 Novel Loci of Urinary Human Metabolic Individuality. PLoS Genet. 11, e1005487 (2015).

70. Trbojević Akmačić, I. et al. High Throughput Glycomics : Optimization of Sample Preparation. Biochem. 80, 934–942 (2015).

71. Lappalainen, T. et al. Transcriptome and genome sequencing uncovers functional variation in humans. Nature 501, 506–11 (2013).

